# Recurrent excitation between motoneurones propagates across segments and is purely glutamatergic

**DOI:** 10.1101/160150

**Authors:** G.S. Bhumbra, M. Beato

## Abstract

Spinal motoneurones constitute the final output for the execution of motor tasks. In addition to innervating muscles, motoneurones project excitatory collateral connections to Renshaw cells and other motoneurones, but the latter have received little attention. We show that motoneurones receive strong synaptic input from other motoneurones throughout development and into maturity with fast type motoneurones systematically receiving greater recurrent excitation than slow type motoneurones. Optical recordings show that activation of motoneurones in one spinal segment can propagate to adjacent segments even in the presence of intact recurrent inhibition. Quite remarkably, while it is known that transmission at the neuromuscular junction is purely cholinergic and Renshaw cells are excited through both acetylcholine and glutamate receptors, here we show that neurotransmission between motoneurones is purely glutamatergic indicating that synaptic transmission systems are differentiated at different post-synaptic targets of motoneurones.

## Introduction

Motoneurones (Mns) are the ultimate neural targets of effector commands issued from the central nervous system. Their activity is modulated by an intricate network of interneurones (Kiehn, 2016) that affect the spatial and temporal distribution of excitation to different motor pools (McCrea and Rybak, 2008). Mns also receive direct inputs from supraspinal tracts and sensory afferents, and their outputs are not confined to the peripheral muscles, but also include excitatory collateral terminals to Renshaw cells (RCs).

Early anatomical studies in the cat have shown that Mn collaterals invade the motor nuclei region and form synaptic contacts with other Mns (Cullheim et al., 1977). Electrophysiological evidence of functional connectivity between Mns has been presented in the adult cat (Gogan et al., 1977), tadpole (Perrins and Roberts, 1995), neonatal (Ichinose and Miyata, 1998) and juvenile (Jiang et al., 1991) rat, and newborn mice (Nishimaru et al., 2005). However despite these initial reports, no study to date has reported systematically the extent, distribution, and pharmacology of direct monosynaptic connections between Mns.

Here we demonstrate strong recurrent excitation between Mns that is maintained throughout development into maturity. Fast type Mns receive greater recurrent excitation than slow type Mns. Under normal physiological conditions, recurrent excitation can override recurrent inhibition and Mn firing in one spinal segment propagates to neighbouring segments. Remarkably, while acetylcholine and a mixture of acetylcholine and glutamate act at the neuromuscular junction and RC synapses respectively (Mentis et al., 2005, Nishimaru et al., 2005), neurotransmission between Mns is purely glutamatergic.

## Results

We performed paired recordings to measure the efficacy of unitary connections in fluorescently labelled Mns innervating gastrocnemius. Simultanous infrared and confocal imaging was used to identify and patch fluorescent Mns in a dorsal horn ablated spinal cord (Figure 1A). Strychnine (0.5 μM) and gabazine (3 μM) were applied to block recurrent inhibition. Mns were patched in whole-cell voltage clamp while putative pre-synaptic cells were stimulated in a loose-cell attached configuration (see Methods) until an evoked response was detected in the post-synaptic cell. Figure 1B shows an example of a paired recording with an average evoked current of *−*34 pA. A location map constructed from 14 out of the 18 recorded pairs (Figure 1C) shows that connected Mns tended to be within 150 μm from one another, but with no systematic relationship between distance and size of response (range 11–125 pA).

**Figure 1:**
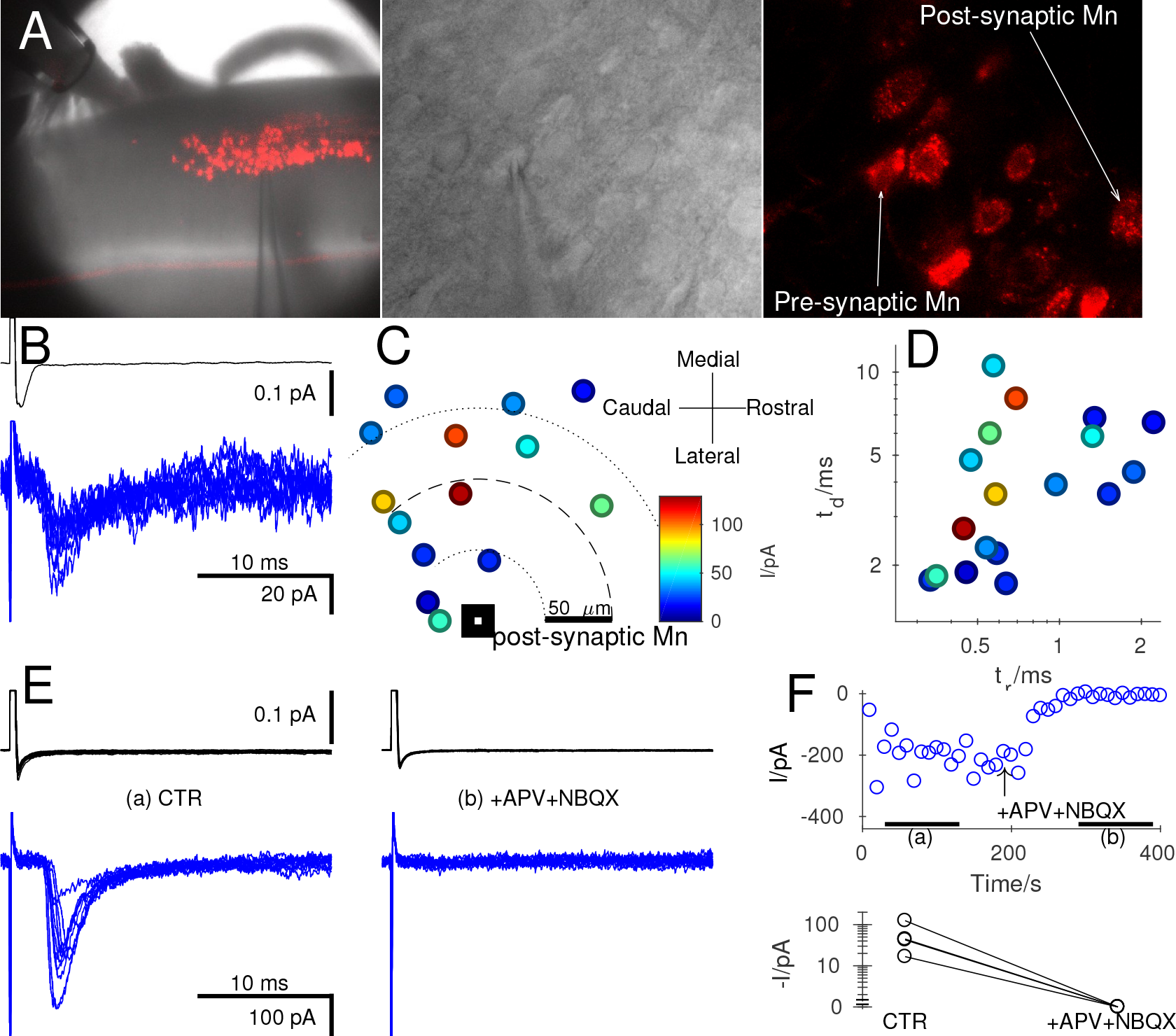
Paired recordings from motoneurones showed small unitary currents that were purely glutamatergic. Prior intramuscular injection of gastrocneumius with CTB-Alexa-Fluor-555 fluorescently labelled the dorsal motor column (A) within the coronal preparation (left) for simultaneous visualisation of motoneurones using intrared (middle) and confocal (right) optics. Pre-synaptic cells were stimulated in loose cell-attached voltage-clamp (B, upper trace) while recording evoked post-synaptic responses in whole-cell voltage clamp (B, lower trace). Connected motoneurones were usually within 150 μm of one other and in most paired recordings (14/18) their respective locations were recorded. Graph C plots the relative position of each pre-synaptic Mn in relation to the post-synaptic cell, with colour-coded size of the corresponding reponses. A similar colour-code and scale is used in graph D, showing decay time (*t_d_*) against the rise time (*t_r_*). Using oblique slice preparations (see text), we investigated the pharmacology of the synapse. Panel E shows a representative recording in control (CTR, left) and following glutamatergic blockade (E, right) using APV and NBQX. The time course of changes in evoked responses during the bath application of the antagonists showed complete suppression of currents (F, top). Similar effects were observed for all four paired recordings (F, bottom).

The rise and decay times of evoked currents (Figure 1D, response size colour coded) were fast, with a median rise time of 0.59 ms and decay time of 3.75 ms. There was however no correlation between either of the kinetic parameters and the size of response. In oblique slice preparations (see Methods) we assessed the pharmacology of evoked responses (Figure 1E). The postsynaptic current was fully abolished by bath-application of 50 μM APV and 2 μM NBQX, to block AMPA and NMDA receptors respectively (Figure 1F, top). Identical results were obtained from all four pairs tested (Figure 1F, bottom).

Since the tested unitary connections might have represented a specific local subset from the entire population of Mn-Mn synapses, we investigated the pharmacology of currents evoked by ventral root (VR) stimulation, thus pooling responses to all inputs from a given segment (Figure 2A). In the example of Figure 2A-C, we simultaneously recorded from a Renshaw cell (RC, Figure 2B, top, red) and a Mn (Figure 2B, bottom, blue). The RC response shown includes a second component originating from a gap junction (Lamotte d’Incamps et al., 2012), contacting a neighbouring RC in which VR stimulation evoked an action potential.

**Figure 2:**
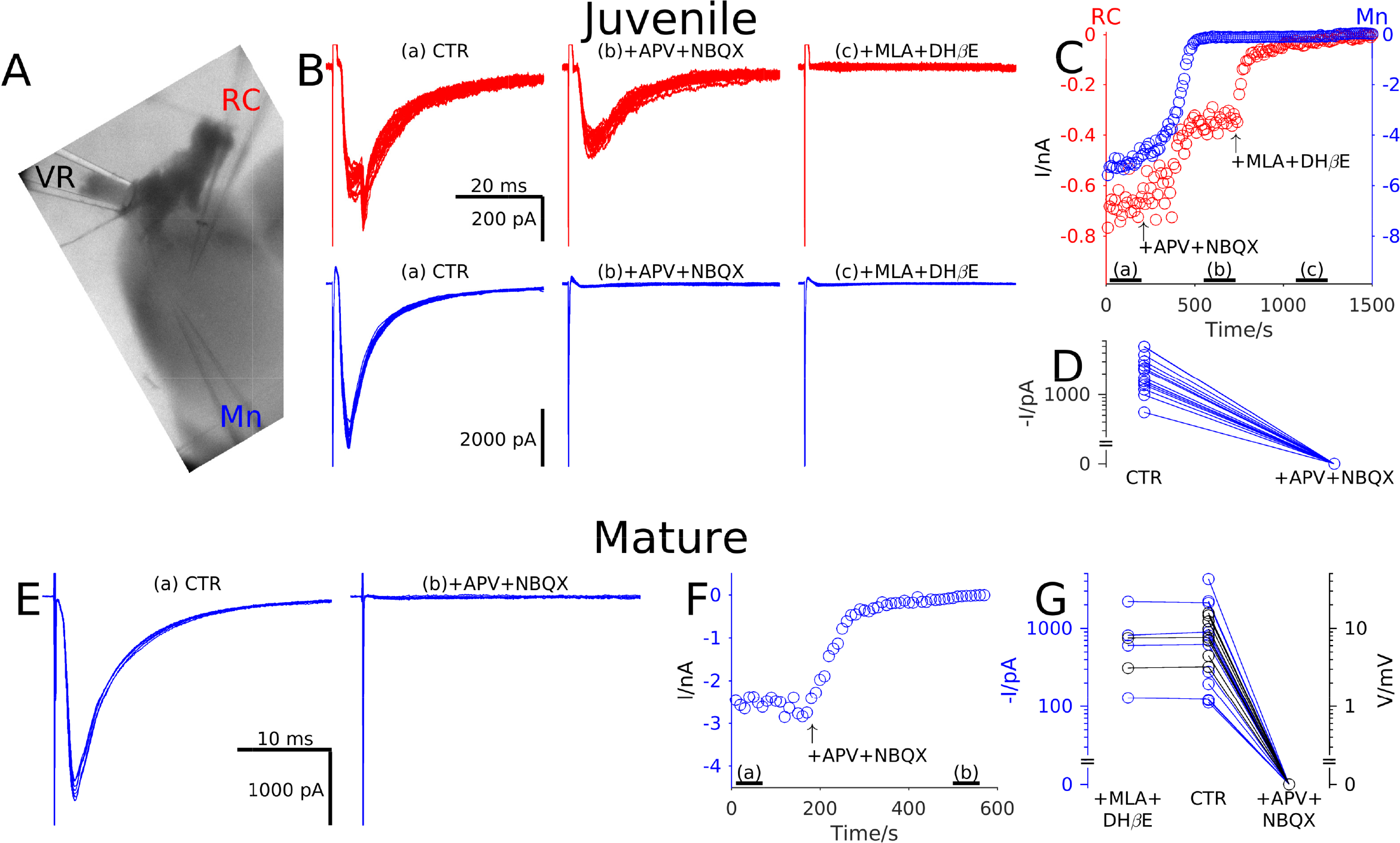
Electrophysiological recordings from both juvenile and mature preparations showed that recurrent excitation in motoneurones is purely glutamatergic. In oblique slice preparations (A) a suction electrode was used to stimulate the ventral root (VR) while recording respones from a motoneurone (Mn) and, in the illustrated example, also from a Renshaw cell (RC). note that the RC response shown includes a second component originating from a gap junction with a neighbouring RC. Panel B illustrates post-synaptic responses (stimulus artefacts truncated) of the RC (red, top) and Mn (blue, bottom) in control (left), in the presence of glutamatergic antagonists (middle) and following block of cholinergic transmission with MLA and DH*β*E (right). The time course of changes in currents (C, top) during application of the antagonists showed partial attenuation of RC responses and full suppression of Mn responses in the presence of glutamatergic antagonists. RC currents were only abolished by block of cholinergic receptors. In all Mns tested the rEPSC was completely abolished by glutamatergic antagonists (D). Similar experiments were performed on mature preparations in which Mn responses to VR stimulation were recorded in control (E, left) and during glutamatergic blockade (E, right). Once again the time course of changes in reponses (F) showed complete suppression of Mn responses in the presence of glutamatergic an-tagonists. Graph G summarises the group data from Mns recorded in voltage clamp (black) and current clamp (blue) showing no effect following bath application of cholinergic antagonists whereas glutamatergic blockade completely abolishes responses.

Whereas bath application of glutamate antagonists resulted in a reduction of the RC response to approximately 50 %, the response in the Mn is completely abolished. The remaining cholinergic component of the response in the RC was blocked by further application of 10 nM MLA and 5 μM DH*β*E to block *α*7 and *αβ* receptors respectively (Figure 2C). Group data from 16 Mn recordings are illustrated in Figure 2D). The mean latency (*±*S.E.M.) of responses of 1.60*±*0.13 ms was consistent with monosynaptic responses to ventral root stimulation. In all cases application of glutamate antagonists resulted in complete suppression of evoked currents (Figure 2D).

While the data from Figure 2D were obtained from juvenile mice (P7-14), we performed similar recordings from more mature animals (P15-25) to determine whether pure glutamatergic transmission is preserved throughout development. Figure 2E-F shows that the response is fully suppressed by glutamatergic blockade. In 20 Mns recorded in voltage-clamp (black, Figure 2G), or in current-clamp (blue, to reduce the duration of the stimulus artefact), glutamatergic antagonists entirely suppressed responses, whereas prior cholinergic blockade had no significant effect (*n* = 6, Wilcoxon’s sign-rank *z* = *−*0.53, *P* = 0.600).

We next investigated whether recurrent excitation could propagate across segments in coronal preparations (see Methods) from juvenile mice (P7-14) in which Mns innervating gastrocnemius were labelled. Figure 3A illustrates recurrent excitatory post-synaptic currents (rEPSCs) recorded in L5 (left) and L4 (right) Mns while stimulating the L5 (upper, blue trace) or L4 (lower, red trace) ventral root (VR). The rEPSC size from 43 recordings from L4 and L5 Mns is plotted against the distance from the L4/L5 border, colour coded to represent responses evoked by L4 (red) or L5 (blue) VR stimulation (Figure 3B). There were no obvious differences in rEPSC size between the two stimulated roots or between L4 and L5 motoneurones. Comparison of rEPSCs from responses to VR stimulation from the same segment or neighbouring segment showed no significant differences (Figure 3B right, Wilcoxon’s rank-sum *z* = 0.61, *P* = 0.541).

**Figure 3:**
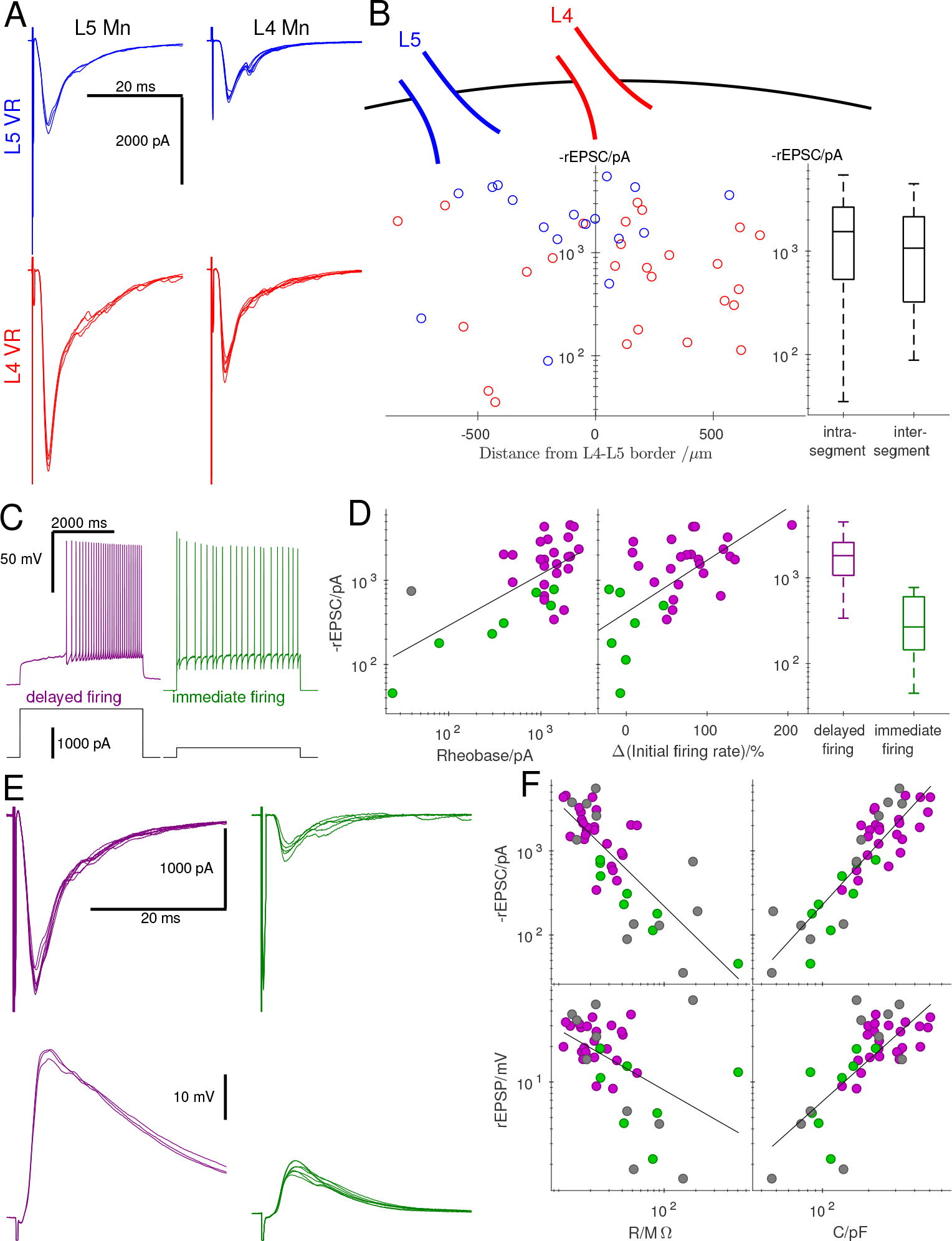
Recordings from coronal preparations showed that the magnitude of rEPSCS was related not to the position of the Mn but to its firing type. Example traces of evoked responses (stimulus artefacts truncated) of motoneurones located in L5 (A, left) and L4 (A, right) segments are illustrated following stimulation of the L5 (A, top, blue) and L4 (A, bottom, red) ventral root; the response in the top right trace exhibited a late component, suggesting activation of a disynaptic pathway. Panel B shows the size of rEPSCs recorded from all Mns against their distance from L4-L5 border (rostral positive) colour-coded according to whether L5 (blue) or L4(red) VR was stimulated. There was no systematic association between rEPSCs and position as shown in the box-and-whiskers plot (B, right) comparing intra-segmental to inter-segmental responses. Two Mn cell types were distinguished using current-clamp recordings on the basis of whether at rheobase, positive current application elicited delayed (C, left, purple) or immediate (C, right, green) firing. Delayed firing cells (purple) were associated with a high rheobase (D, left), an accelerating initial firing rate (D, middle), and large evoked rEPSCs (D, right) in comparison to immediate firing cells (green). The traces in E illustrate representative responses to VR stimulation from a delayed firing cell (purple, left) and immediate firing cell (green, right) recorded in voltage-clamp (top) and current-clamp (bottom). Graph F shows the group data, plotting the rEPSCs and rEPSPs against cell resistance and capacitance, using grey circles to denote cells that were not identified by their firing pattern at rheobase. In all four cases, correlations were observed demonstrating that responses were greater in larger Mns, which tended to be of the delayed firing type.

The mean latency (*±*S.E.M.) of responses of 1.64*±*0.09 ms was consistent with monosynaptic activation and not significantly different from those from oblique slices (1.60*±*0.13 ms, Wilcoxon’s rank-sum *z* = *−*0.90, *P* = 0.384). Latencies of responses from Mns in neighbouring segments (1.77*±*0.13 ms) were longer compared to those recorded from within the same segment that was stimulated (1.43*±*0.09 ms, Wilcoxon’s rank-sum *z* = *−*2.43, *P* = 0.015). The corresponding response jitter, quantified using the standard deviation of the latencies, were very small both within (0.08*±*0.03 ms) and across (0.08*±*0.01 ms) segments and a comparison between the two showed no significant difference in jitters (Wilcoxon’s rank-sum *z* = *−*0.58, *P* = 0.559). These results demonstrate that while the latency of synaptic responses may be greater across segments compared to within segments, they are both mediated by monosynaptic connections.

We then assessed whether the magnitude of recurrent excitation was related to the intrinsic properties of post-synaptic Mns. Two types of Mns were identified according to their firing pattern at rheobase in current clamp recordings (Leroy et al., 2014). The first type (Figure 3C, left, purple) has high a rheobase and produces *delayed firing* with a pronounced increase in firing rate during positive current application and is associated with fast-type units. By contrast the second type (Figure 3C, right, green) has a lower rheobase and *immediate firing* with little change in spike frequency, characteristic of slow-type units (Leroy et al., 2014). High rheobase (Figure 3D, left) and accelerating initial firing (Figure 3D, middle) were correlated with the size of rEPSCs, with median values of 1814 pA in the delayed firing cells and 267 pA in the immediate firing cells. Comparison between the two groups confirmed a significant difference (Figure 3D right, Wilcoxon’s rank-sum *z* = 3.72, *P <* 0.001).

Differences between the two cell types are also associated with their passive properties, with delayed firing Mns showing lower resistances (median 22 MΩ) and higher capacitances (median 237 pF) than their immediate firing counterparts (median resistance 44 MΩ, median capacitance 125 pF) both at statistically significant levels (Wilcoxon’s rank-sum *|z| ≥* 2.94, *P ≤* 0.003). Recurrent excitatory responses were recorded from delayed firing (Figure 3E, purple) and immediate firing (Figure 3E, green) cells in both voltage clamp (top) and cur-5 rent clamp (bottom). Pooling all cell types together, correlations were observed between the size of response and resistance or capacitance (Spearman’s *|r| ≥* 0.516, *P <* 0.001, Figure 3F).

The presence of strychnine and gabazine during electrophysiological recordings precluded evaluation of whether recurrent excitation could override recurrent inhibition, We therefore conducted calcium imaging experiments in mice selectively expressing GCaMP6s in Mns to evaluate the propagation of recurrent excitation across different segments with recurrent inhibition intact. Figure 4A-C illustrates a coronal preparation with a suction electrode applied to the L5 ventral root (Figure 4A, left). Calcium signals were acquired throughout the dorsal motor column of L4 and L5 (Figure 4A, middle) with 146 ms frame interval before, during, and following a train of three VR stimulations at 30 Hz. The signal from regions of interest, defined within the outline of Mn somata, was evaluated for the period of acquisition under control conditions, in the presence of 0.5 μM strychnine and 3 μM gabazine, and following application of 50 μM APV and 2 μM NBQX (Figure 4A, right).

**Figure 4:**
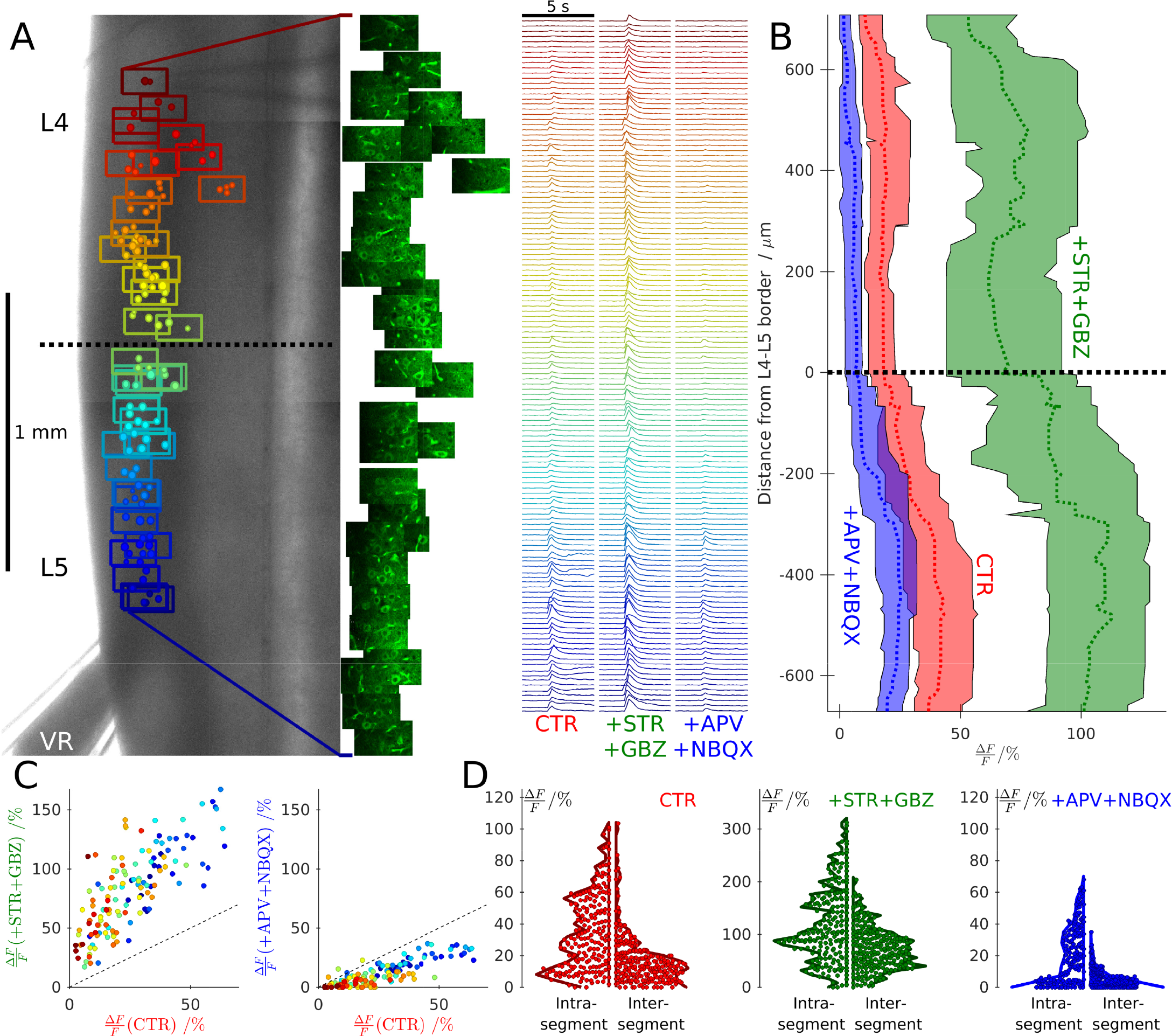
Calcium imaging from coronal preparations showed that recurrent excitation from one segment can evoke spikes in adjacent segments with intact recurrent inhibition, but these responses are abolished by glutamatergic antagonists. An example of such a recording is illstrated in A, which overlays the Mn positions within optical fields colour-coded by position, on top of a low-magnification image (A, left) of a coronal L4-5 preparation from a ChAT-GCaMP6s mouse in which Mns express GCaMP6s. A suction pipette was used to stimulate the L5 ventral root (VR) while acquiring fluorescence intensities throughout each field (A, middle). Changes in Mn fluorescence were measured and plotted against time (A, right) under control conditions (CTR) in the presence of pharmacological blockade of inhibition (+STR+GBZ) and addition of further antagonists to block glutamatergic neurotransmission (+APV+NBQX). The results for this experiment are summarised in graph B, in which running medians and inter-quartile ranges of responses are plotted for the three conditions against the distance from the L4-5 border (rostral positive). Recurrent excitation from L5 evoked firing in Mns throughout the L4 segment and these responses were enhanced by inhibitory antagonists and abolished by glutamatergic blockade. These effects are illustrated by the graphs in C which plots the signal response under control conditions with that in the presence of antagonists of inhibition (C, left) and glutamatergic bockade (C, right), preserving the colourcoding of position in the rostral-caudal axis used in panel A. Violin plots summarise the group data, comparing intra-segmental and inter-segmental responses, showing the distribution of the magnitude of responses from all Mns under control conditions (D, left), inhibitory blockade (E, middle; on a different y-scale), and in the additional presence of glutamatergic antagonists (D, right).

In control, recurrent excitation evoked spikes in Mns from both L4 and L5 segments, as shown by running the medians and inter-quartile ranges of the relative fluorescence signal (Figure 4B, red) throughout both segments. Bath application of strychnine and gabazine resulted in substantial amplification of responses throughout the motor column (Figure 4B, green) whereas additional application of glutamatergic antagonists abolished responses from the L4 segment and substantially attenutated those from L5 Mns (Figure 4B, blue). The residual response in L5 Mns reflects antidromic activation. Scattergrams, colour-coded by regions, comparing control responses to those during application of inhibitory antagonists (Figure 4C, left) and additional glutamatergic blockade (Figure 4C, right) confirm that while responses were greater in the lumbar regions closer to the stimulated VR, the relative effects of block of recurrent inhibition, or excitation, were similar throughout L4 and L5.

Group data from 461 Mns from 7 preparations are shown in Figure 4D, comparing responses within and across segments for the three conditions. In control, responses were significantly greater within the stimulated segment compared to outside (Figure 4D, left, Wilcoxon’s rank-sum *z* = 8.50, *P <* 0.001) and these difference were maintained after block of inhibition 6 (Figure 4D, middle) and excitation (Figure 4D, right) (*z ≥* 7.23, *P <* 0.001). Pooling Mns from both segments, blockade of inhibition consistently increased the signal (Wilcoxon’s sign-rank *z* = *−*18.59, *P <* 0.001) whereas a significant reduction in signal was observed following additional application of glutamatergic antagonists (*z* = 18.20, *P <* 0.001). Residual firing was mostly confined to Mns within the stimulated segment through antidromic activation, thus confirming the purely glutamatergic nature of recurrent excitation.

## Discussion

Our experiments show that strong recurrent excitation between Mns is maintained throughout development and fast Mns receive greater recurrent excitation than slow ones, We demontrate that synaptic transmission between Mns is purely glutamatergic. While it could be argued that the observed small unitary post-synaptic responses (*∼*100 pA) would have little effect on the excitability of Mns whose somata are very large, ventral root stimulation evoked responses usually exceeding 1 nA indicating extensive convergence of segmental Mn populations.

The variation in the magnitude of rEPSCs is associated with Mn classification into delayed and immediate firing types. Larger responses were systematically observed in the delayed firing, low resistance, and high capacitance cells. Our results are thus consistent with a structural connectivity in which the fast-type larger Mns receive stronger recurrent excitation compared to slow-type smaller cells. This pattern of connectivity suggests that recurrent excitation could play a role in sequential recruitment of fast-type units during motor tasks in which progressively increasing muscular forces are needed. Alternatively, recurrent excitation might represent a closed-loop amplification circuit that reinforces and increases the firing rate preferentially in fast-type Mns and thus rapidly increase muscle contraction strength when required.

In neonatal animals, VR stimulation can induce fictive locomotion (Mentis et al., 2005) and entrain the spontaneous rhythmic bursting induced by block of inhibition (Bonnot et al., 2009). Furthermore, optogenetic activation or silencing of motor pools alters the frequency and phase of chemically induced fictive locomotion (Falgairolle et al., 2017). These effects cannot be explained solely by recurrent excitation and may provide evidence for Mn collaterals contacting unidentified interneurones (Machacek and Hochman, 2006). In ourelectrophysiological recordings during pharmacological blockade of recurrent inhibition, we often observed a late disynaptic component that may result from orthodromic activation of Mn pools not antidromically activated by VR stimulation. Such recruitment implies the existence of a positive-feedback amplifying circuit whose tendency to reverberate may be suppressed by recurrent inhibition.

However, the recurrent excitation characterised in the present study comprises predominantly a monosynaptic component and this is evidenced by three observations. First, the connectivity between Mn pairs must have been monosynaptic. Second, the latency of responses within (1.43*±*0.09 ms) and between (1.77*±*0.13 ms) lumbar segments were within the time-scale of neurotransmission through only a single synapse. Finally, the response jitters within (0.08*±*0.03 ms) and between (0.08*±*0.01 ms) segments were very small and virtually identical. These observations are only consistent with a monosynaptic connectivity between Mns both within the same segment and across neighbouring segments. While the occurrence of synaptic projections between Mns crossing spinal segments may be regarded as unusual, it is perfectly compatable with the known rostro-caudal distribution of Mn dendritic trees which may span over a millimetre in juvenile mice with little or no change into adulthood (Li et al., 2005).

A glutamate receptor-dependent effect on Mn EPSPs evoked by VR stimulation has been reported previously (Jiang et al., 1991) but it was attributed to afferent fibres within the root (Coggeshall, 1980), a possibility now excluded by subsequent labelling studies (Mentis et al., 2005). A previous study has reported a purely cholinergic response to VR stimulation in a small proportion (2/9) of Mns (Nishimaru et al., 2005). Across all electrophysiological recordings of the present study however, there was not a single instance of a cholinergic component. The origin of such a discrepancy may result from differences in maturity, since in the previous study (Nishimaru et al., 2005) neonatal mice (P0-P4) were used, while our experiments were performed on mice of at least weight bearing age (P7-P25).

Neurotransmission between Mns is purely glutamatergic, yet following normal maturation, the neuromuscular junction is solely cholinergic (Borodinsky and Spitzer, 2007) and synaptic transmission of recurrent collaterals onto Renshaw cells is mixed with both cholinergic and glutamatergic components (Lamotte d’Incamps et al., 2017). This remarkable dissociation demonstrates a differentiation of neurotransmission systems on the basis of the different post-synaptic targets of Mns. However, the presence of vesicular glutamate transporters in Mn collateral terminals is still controversial. Immunohistochemistry and *in situ* hybridization studies have reported the expression of the vesicular transporter VGlut2 in some Mns terminals onto RCs that are either positive (Nishimaru et al., 2005) or negative (Herzog et al., 2004) for the vesicular acetylcholine transporter. These respective findings indicate either coexistence or segregration of cholinergic and glutamatergic transmission of Mns onto RCs.

Others however have not detected the presence of VGlut2, or any other vesicular glutamate transporter, in Mn terminals (Mentis et al., 2005, Liu et al., 2009). It is possible that such discrepencies arise from undetectable albeit functional expression levels of VGlut2. Another possibility is the existence of an unidentified vesicular glutamate transporter (Mentis et al., 2005, Liu et al., 2009). This hypothesis is supported by the presence of glutamate releasing C-fibres in the dorsal horn that are nevertheless negative for all known vesicular glutamate transporters (Todd et al., 2003, Alvarez et al., 2004). Since many Mn terminals may contain more aspartate than glutamate (Richards et al., 2014), it has been proposed that the released neurotransmitter could be aspartate. However, aspartate alone cannot activate AMPA receptors that mediate responses of RCs (Lamotte d’Incamps and Ascher, 2008) or of the Mns characterised in the present study. Glutamate thus remains the most likely candidate.

## Materials and Methods

All experiments were carried out in accordance with the Animal (Scientific Procedures) Act (Home Office, UK, 1986) and were approved by the UCL Ethical Committee, under project licence number 70/7621. Experiments were performed on preparations obtained from male or female mice bred using a C57BL/6J background. For electrophysiological experiments with simultaneous recordings from motoneurones (Mns) and Renshaw cells (RCs) a transgenic strain, in which the enhanced green fluorescent protein (EGFP) is expressed under the control of the promotor of the neuronal glycine transporter GlyT-2 (Zeilhofer et al., 2005), was used to label glycinergic interneurones.

### Spinal cord preparations

Following anaesthesia by intraperitoneal injection of a mixture of ketamine/xylazine (80 mg/kg and 10 mg/kg respectively), both juvenile and mature mice were decapitated and the spinal cord dissected in normal ice cold aCSF containing (in mM) 113 NaCl, 3 KCl, 25 NaHCO_3_, 1 NaH_2_PO_4_, 2 CaCl_2_, 2 MgCl_2_, and 11 D-glucose (same solution was used for recording). The spinal cord was then glued onto an agar block and affixed to the chamber of a vibrating slicer (HM 650V, Microm). We used a slicing solution containing (in mM) 130 K-gluconate, 15 KCl, 0.05 EGTA, 20 HEPES, 25 D-glucose, 3 kynurenic acid and ph 7.4 with NaOH (Dugue et al., 2005). For cutting oblique slices, the cord was glued to an agar block cut at a 45 degrees angle, with the ventral side facing the direction of the blade (Lamotte d’Incamps et al., 2017).

For coronally sliced preparations in which the dorsal horns were ablated, the cord was glued horizontally with the ventral surface facing upwards. A blade was used to transect the cord at the L1-L2 boundary at an angle that allowed visualization of the exact position of the central canal under a dissection microscope. The vibratome blade was then aligned to the central canal and the ventral portion of the cord was sliced away from the dorsal part. Alignment with the central canal was essential to ensure a consistent dorsoventral level of the ablation across different preparations and to retain the dorsal motor nuclei near the cut surface of the tissue.

Identical procedures were used for juvenile (P7-14) and mature (P15-25) animals. For older animals we routinely cut the first slice within 8 minutes following decapitation. Since spinal cord preparations are extremely sensitive to anoxia especially prior to slicing, we found that minimizing the time to obtain the first slice consistently resulted in viable preparations with healthy motoneurones (Lamotte d’Incamps et al., 2017).

### Electrophysiology

All recordings from post-synaptic motoneurones were performed with a Molecular Devices Axopatch 200B amplifier, filtered at 5 kHz and digitized at 50 kHz. Patch pipettes were pulled to resistances in the range of 0.8–2 MΩ when filled with (in mM) 125 K-gluconate, 6 KCl, 10 HEPES, 0.1 EGTA, 2 Mg-ATP, pH 7.3 with KOH, and osmolarity of 290–310 mOsM. During voltage-clamp recordings, Mns were clamped at *−*60 mV with series resistances in the range of 2–10 MΩ compensated by 60-80%.

During paired recordings, loose cell attached stimulation was used to evoke spikes in puta-tive pre-synaptic motoneurones using an ELC-03X (NPI Instruments) amplifier and a 4–5 MΩ pipette filled with normal aCSF (Bhumbra et al., 2014). Ventral root stimulation was delivered to evoke recurrent excitatory post-synaptic currents in motoneurones using a glass suction electrode whose tip was cut to correspond with the size of the ventral root (Moore et al., 2015). The stimulation intensity was increased until the size of the rEPSC remained constant, typically at 5*×* threshold. In order to exclude direct stimulation of the ventral white matter, ventral roots were only used if they were of sufficient length to afford no possible physical contact between the slice and suction pipette. This was tested before and after each recording by confirming that side-to-side movement of the suction pipette resulted only in movement of the root and not the slice.

For measuring the size of the excitatory response in some Mns, where it was necessary to prevent action potentials, cells were hyperpolarised below their resting membrane potential. Measurements of synaptic current and potentials from these recordings were adjusted to their predicted value at *−*60 mV assuming a reversal potential of 0 mV for excitatory conductances. All electrophysiological experiments were performed in the presence of 0.5 μM strychnine and 3 μM gabazine. Where indicated, excitatory receptors were blocked using D-2-amino-5-phosphonopentanoic acid (APV), 1,2,3,4-tetrahydrobenzo(f)quinoxaline-7-sulphonamide (NBQX), methyllycaconitine (MLA) or dihydro-*β*-erythroidine (DH*β*E).

### Intramuscular injections

In order to label motoneurones innervating the ankle flexor gastrocnemius muscle, intramuscular injections were performed 2-5 days prior to recording. Inhalant isofluorane was used for the induction and maintenance of anaesthesia. Traction was applied to the lower limb and an incision was made through the skin and deep fascia overlying the muscle. A Hamilton syringe loaded with a glass needle was used to inject 1 μl of CTB-Alexa-Fluor-555 (0.2% in 1*×* phosphate buffer saline) into the middle of the muscle belly over a period of at least 1 minute. The skin was closed by suture using a buried stitch before cessation of anaesthesia and recovery.

## Calcium imaging

Calcium imaging experiments were performed on animals selectively expressing the genetically encoded calcium indicator GCaMP6s in motoneurones. These mice were generated by crossing mice expressing Cre under the control of choline-acetyltransferase (ChAT-Cre, JAX mouse line number 006410) with animals with the gene expressing GCaMP6 flanked by a flox-Stop cassette (JAX mouse line number 028866, Madisen et al., 2015). Upon recombination with Cre, GCaMP6 is selectively expressed in motoneurones and other cholinergic cells of the offspring. Since the only other population of ChAT positive lumbar spinal cells are the cholinergic partition neurones located around the central canal, there was no ambiguity in the identification of motoneurones from their basal GCaMP6 fluorescence and position within the motor nuclei.

Dorsal horn ablated coronal slice preparations (P9-12) were used for imaging experiments to visualise the dorsal motor nuclei in the L4 and L5 segments containing Mns innervating tibialis anterior, gastrocnemius, and peroneus longus close the cut surface. A laser scanning confocal unit (D-Eclipse C1, Nikon) with a diode laser (*λ* =488 nm, power output from optic fibre 3–5 mW) was used to locate and record calcium signals from Mns reaching a depth of approximately 100 μm from the surface. Fields of 128x64 pixels (pixel size 1.38 μm and dwell μs) were scanned with a frame interval of 146 ms over different regions throughout the dorsal motor column. Trains of 3 stimulii at 30 Hz were delivered to the ventral root (L4 or L5) while images were being acquired from at least 1 s before the onset of the first stimulus pulse. For each field, calcium signals were acquired for a total of 35 frames corresponding to approximately 5 s, and the position of each field was recorded.

Post-hoc analysis was performed to quantify Mn responses. Within each field, single Mns were identified by their fluorescence and regions of interests were defined by the contour profiles of their somata. The time course of excitation was measured using the change in mean fluoresence following stimulation divided by the baseline average. In some cases, slow drifts in fluorescence was corrected by fitting an exponential to the initial trace before the stimulus. Changes in fluorescence exceeding two standard deviations of the baseline noise were measured over a 1 s window following ventral root stimulation.

## Acknowledgments

This work was supported by grants from the Leverhulme Trust (grant number RPG-2013-176) and the Biotechnology and Biological Sciences Research Council (BBSRC, grant number BB/L001454) to MB. We are grateful to Professor John Wood for providing the GCaMP6s mice and to Professor Rob Brownstone for helpful discussions during the course of this study.

